# Theoretical investigation of stochastic clearance of bacteria: First-passage analysis

**DOI:** 10.1101/446591

**Authors:** Hamid Teimouri, Anatoly B. Kolomeisky

## Abstract

Understanding mechanisms of bacterial eradication is critically important for overcoming failures of antibiotic treatments. Current studies suggest that the clearance of large bacterial populations proceeds deterministically, while for smaller populations the stochastic effects become more relevant. Here we develop a theoretical approach to investigate the bacterial population dynamics under the effect of antibiotic drugs using a method of first-passage processes. It allows us to explicitly evaluate the most important characteristics of the bacterial clearance dynamics such as extinction probabilities and extinction times. The new meaning of minimal inhibitory concentrations for stochastic clearance of bacterial populations is also discussed. In addition, we investigate the effect of fluctuations in the population growth rates on dynamics of bacterial eradication. It is found that extinction probabilities and extinction times generally do not correlate with each other when random fluctuations in the growth rates are taking place. Unexpectedly, for a significant range of parameters the extinction times increase due to these fluctuations, indicating a slowing in the bacterial clearance dynamics. It is argued that this might be one of the initial steps in the pathway for the development of antibiotic resistance. Furthermore, it is suggested that extinction times is a convenient measure of bacterial tolerance.

## Introduction

The rise of pathogenic bacteria that are resistant to antibiotics is one of the major global health threats of the 21st century. High mortality rates and increasing health care costs associated with fighting the bacterial infections call for designing new effective therapeutic strategies [7, 26]. A major challenge in overcoming treatment failures is coming from ineffective eradication of antibiotic-susceptible bacteria [12, 29, 34]. Despite the introduction and wide application of a very large range of antibiotics since the 1940s, important aspects of how antibiotics clear bacterial population at all levels (molecular, cellular and population) remain not well clarified. A deeper understanding of the underlying dynamics of bacterial clearance requires not only extensive laboratory studies, but also a development of new theoretical approaches to investigate the bacterial response to antibiotics [2].

Majority of current experimental and theoretical studies focus on the eradication of initially large quantities of bacteria [14, 25, 28], and it was shown that a deterministic picture describes well the decrease in these bacterial populations [10, 28]. In this deterministic framework, the dynamics of bacterial population exposed to antibiotic is characterized by a minimum inhibitory concentration (MIC), the minimal drug concentration required to inhibit bacterial growth [10, 13, 28]. The MIC can be regarded as a threshold on the antibiotic concentration such that only above MIC a bacterial population can undergo full extinction, while for concentrations below MIC the infection will never disappear.

However, it can be argued that it is also critically important to investigate the clearance dynamics for small bacterial populations. Failure to completely eradicate a population of bacteria can have two main consequences. First, even a small number of surviving bacteria can restore infections [19]. Second, certain strains of surviving cells may develop antibiotic resistance, which, in turn, can complicate subsequent therapies [11, 17, 20]. Therefore, the effective treatment of infections requires not only reducing a large population number to a small number, but also the complete eradication of the bacterial population [4, 32, 35].

Despite earlier technical problems [14, 25], recent experiments were able to quantitatively investigate the antibiotic-induced clearance of small bacterial populations [9]. It was demonstrated that stochastic factors play much more important roles at these conditions. For example, Coates et al showed that even in sub-MIC antibiotic concentration, bacterial population decline with non-zero probability [9]. This means that under the same conditions some populations experience growth with cells continuously dividing, while other populations quickly extinct. A Markovian probabilistic birth-and-death model was introduced to uncover the relationship between the extinction probability and the antibiotic concentrations [9]. This stochastic approach predicted that antibiotics induce fluctuations in bacterial population numbers. These fluctuations, in turn, lead to stochastic nature of the clearance of small bacterial populations.

Although the Markovian model developed by Coates et al successfully described the experimental observations, it cannot predict an extinction time, i.e., the mean time at which the given number of bacterial cells will go to zero. This is a very important property of the bacterial population clearance dynamics because it gives a better measure of the efficiency of the antibiotic treatments than the extinction probability. One could use an analogy with thermodynamic and kinetic descriptions of chemical processes. Thermodynamics gives the probability for the process to happen, but if the process is actually taking place in real times is determined by kinetic rates. In our language, this means that the large extinction probability might not always correlate with fast removal of bacterial infection. While the extinction probability can give a qualitative measure of the bacterial population dynamics, the extinction time is much more useful in quantitative characterization of the bacterial resistance and tolerance. It seems that the development of new drugs and new therapies in fighting against bacteria should utilize this quantity as a measure of their success.

In this study, we developed a discrete-state stochastic model of the antibiotic-induced clearance of bacteria that employs a method of first-passage probabilities. This method has been successfully utilized to analyze multiple processes in Chemistry, Physics and Biology [21, 27, 33]. It allows us to quantitatively describe the stochastic dynamics of bacterial eradication by explicitly calculating extinction probabilities and extinction times and clarifying the physical meaning of MIC. Our method is also applied to investigate the effect of fluctuations in the growth rates on the stochastic clearance of bacterial populations. These fluctuations can be attributed to various environmental factors such as availability of nutrients, changes in osmolarity and other factors [30]. Our results suggest that these fluctuations influence the extinction probabilities and extinction times differently. There is a large range of antibiotic concentrations when the extinction times increase due to the fluctuations, and this corresponds to the slowdown of the dynamics of bacterial eradication. We speculate that this might be a first step in the developing of antibiotic resistance. It is also argued that extinction times is a convenient new measure of bacterial tolerance.

## 1 Model

### 1.0.1 Stochastic clearance with a constant growth rate

We start our analysis by considering a simple stochastic model for the clearance of bacteria as shown in figure 1. Our goal is to obtain a minimal theoretical description of the bacterial clearance dynamics. For this reason, the model is characterized by only two parameters: the rate of cell growth λ and the rate of cell death *ϕ*: see figure 1. The bacterial growth rate is generally controlled by the environmental factors such as the availability of nutrients, temperature, osmotic pressure and other factors [30]. When exposed to antibiotics, the cell growth rate can also depend on the antibiotic concentration [16]. For the sake of simplicity, we assume that the cell growth rate is independent of antibiotic concentration and remains constant over different generations, while the cell death rate, *ϕ*, is controlled by the antibiotic concentration. It is also assumed here that if the bacterial population reaches the size *N* the organism hosting the bacteria will die from the infection. This is known as a fixation.

**Figure 1.**
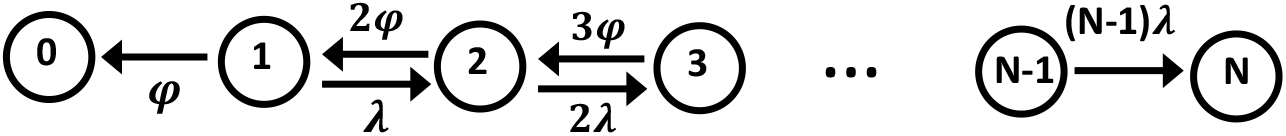
Schematic representation of the single growth-rate model for the clearance of bacteria. Each state *n* (*n* = 0, 1, …, *N*) represents a bacterial population with *n* cells. The states 0 and *N* correspond to the bacterial eradication (no cells in the system), and the fixation (death of the organism), respectively. From each state *n*, the bacterial population can change to the state *n* + 1 (growth) with a total rate *n*λ, or it can jump to the state *n* − 1 (shrinking) with a total rate *nϕ*.

To describe dynamical transitions in the system, we define *F_n_*(*t*) as a probability density function to clear the system from infection at time *t* if the initial population number (so-called inoculum size) is equal to *n* (1 ≤ *n* ≤ *N* − 1). The temporal evolution of this probability function is governed by the following backward master equation [21, 27]:

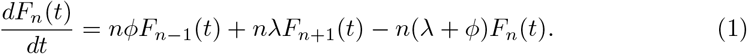

Introducing the Laplace transform of this function, 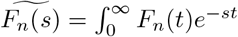, we transform the backward master equation into

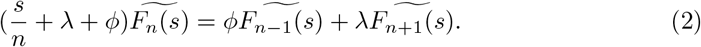

Because we are mostly interested in the stationary dynamic behavior at long times (*s* → 0), the following expansion can be written:

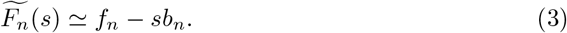

Then 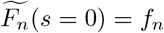 yields the first-passage probability of the bacterial clearance or simply the extinction probability for the bacterial population with the inoculum size *n*. It can be shown that the extinction probability is given by (see appendix I for details)

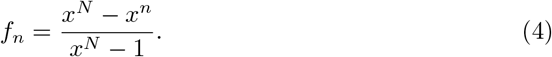

where a parameter *x* = *ϕ*/λ can be viewed as an effective death rate for the bacterial population normalized over the growth rate. Since in our model it is assumed that the growth rate does not depend on the death rate, the extinction probability is determined only by the ratio of *ϕ* and λ.

Our analytical results for the extinction probability are presented in figure 2. The dependence of the bacterial clearance probability [from equation (4)] on the initial size of the bacterial population is given in figure 2(a) for three different values of *x*. For *x* = 1, which in the deterministic picture of bacterial clearance is described as MIC, the extinction probability linearly decreases with the inoculum size, 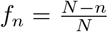. In this case, the growth and the death rates are the same, and the probability of bacterial clearance is proportional to the relative distance from the initial state *n* to the fixation state *N*. The smaller the inoculum size, the larger the probability to eradicate the infection. But even for *n* = 1, the extinction probability is not equal to one 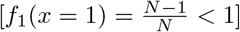. For *x* < 1 (sub-MIC conditions), the extinction probability is a decaying function of the inoculum size *n*. In this case, the growth rate is faster than the death rate, and the larger the inoculum size, the harder for the system to reach the total eradication of the infection (*n* = 0 state). One could also see this more clearly in the limit of *x* → 0 and *N* → ∞ when we have *f_n_* ≃ *x^n^*. This implies that even for sub-MIC conditions (low antibiotic concentrations) the extinction probability is never equal to zero, which is a clear signature of the stochastic effects in the bacterial clearance dynamics. The situation is different for *x* > 1 (large antibiotic concentrations), when the extinction probability is always close to one except in the region near the death state *N*. This can be also seen from the case of *x* ≫ 1 and *N* → ∞ when we obtain *f_n_* ≃ 1 − *x^n−N^*. This result suggests that even for concentrations above MIC the extinction probability is never equal to one, which is again due to the stochastic fluctuations in the system. Our analytical calculations were verified with Monte Carlo computer simulations, in which we utilized typical growth rates associated with bacteria *E. coli*, in range from 1/300 *min*^−1^ to 1/20 *min*^−1^ [30].

**Figure 2.**
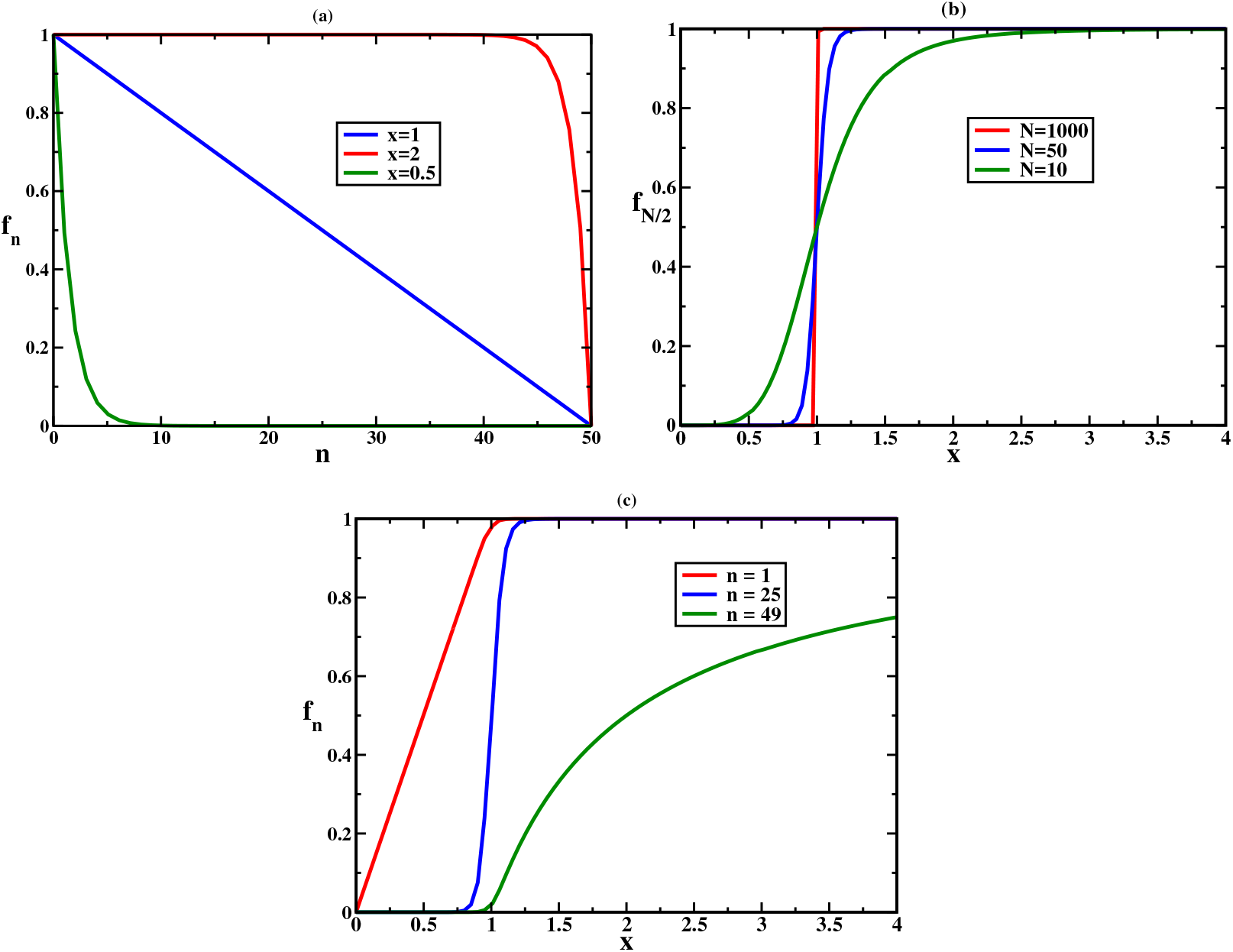
Analytical calculations of extinction probabilities: (a) as a function of the inoculum size for three different values of *x* and *N* = 50; (b) for a specific mid-size inoculum (*n* = *N*/2) as a function of the parameter *x* (*x* = *ϕ*/λ) for three different values of *N*; and (c) as a function of the parameter *x* for three different values of the inoculum size with *N* = 50.

The stochastic effects of the bacterial clearance can be understood better if we consider the extinction probability of a specific inoculum size (*n* = *N*/2), equally distant from the state *n* = 0 (eradication) and *n* = *N* (death), which is plotted in figure 2(b). One can see that the dependence of the extinction probability on *x* follows a logistic sigmoid curve. The steepness of the curve at midpoint (*x* = 1) is controlled by the values of *n* and *N*. In other words, for *x* < 1 the extinction probability is still non-zero while for *x* > 1 it is still less than one. Therefore *x* = 1 does not satisfy the classical definition of MIC as the minimum inhibitory concentration required for clearance. Thus, we need to calculate the effective saturation value for which the extinction probability becomes very high and realistically not much different from one. This might be viewed as an effective MIC for stochastic bacterial clearance. This saturation point is given by(see appendix II)

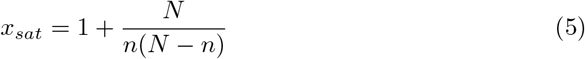

For the special case 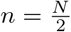, this equation yields 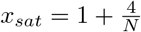. Therefore, as N increases the steepness of curve becomes sharper, such that the extinction probability becomes insensitive to population number while it is ULTRASENSITIVE with respect to *x*. In this case, large population alleviates the stochastic effects in the bacterial clearance, and *x* = 1 yields the minimum inhibitory concentration, as expected.

Theoretical calculations also predict that the extinction probability strongly depends on the inoculum size and on its relative distance to the death state *N*, as illustrated in figure 2c. For *n* = 1 the dependence on *x* is linear for small antibiotic concentrations (*x* < 1), while *n* = *N* − 1 is almost zero for *x* < 1 and it is slowly approaching to one for larger antibiotic concentrations. These different behaviors are again a consequence of the stochastic nature of the bacterial population clearance. A critically important property of the bacterial eradication is how long does it take to clear the host from the infection, which is known as the extinction time. This time scale is crucial for development of new therapies and it can be also useful in quantifying the bacterial tolerance, which is the ability of a bacterial population to survive at longer periods of time exposed to antibiotics [5]. Our first-passage probabilities method is a powerful tool to evaluate this quantity. We define *T_n_* as a mean first-passage time to reach the extinction state (*n* = 0) from the inoculum of size *n*, and this is exactly the extinction time. Using the probability density function *F_n_*(*t*), it can be written as

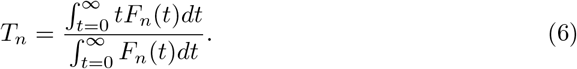

Using the Laplace transform and equation (3), we obtain

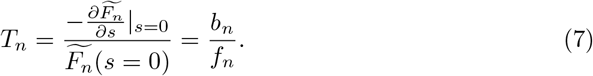

As explained in the appendix I, the extinction time is explicitly given by:

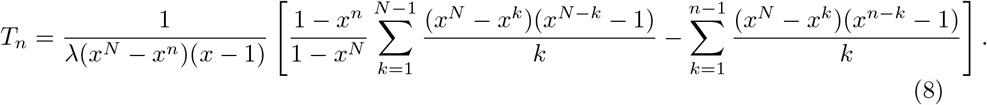

It can be shown that for *x* = 1 the expression for the extinction time takes the form:

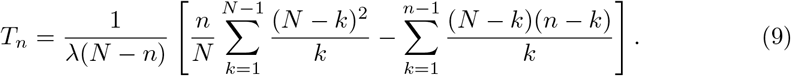

For *x* > 1 and *N* → ∞ the extinction times are given by (see the appendix I),

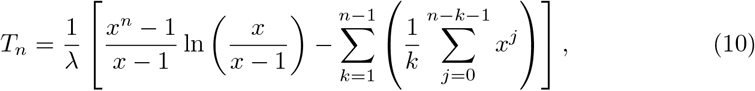

while for *x* → 0 we have

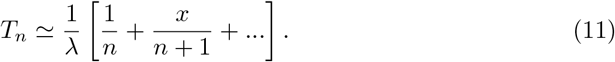

The results of our calculations for the extinction times are presented in figure 3. As expected, it takes longer to clear the infection for larger inoculum sizes (figure 3a). For large antibiotic concentrations (*x* > 1), the extinction time is shorter and it depends weaker on the inoculum size *n*. For small antibiotic concentrations (*x* < 1), the time to eradicate the infection is larger and it is more sensitive to the inoculum size. More interesting behavior is observed when we analyze the extinction time for different antibiotic concentrations: see figure 3b. A non-monotonic behavior as a function of *x* is predicted, and the largest extinction time is observed for MIC conditions (*x* = 1). Increasing the antibiotic concentrations (*x* > 1) shortens the time for bacterial clearance because the drive to infection eradication becomes stronger. However, the surprising observation is that lowering the antibiotic concentrations below MIC (*x* < 1) can also accelerate the bacterial clearance despite the fact that the probability of clearance decreases. This can be explained by the following arguments. In these conditions, only those bacterial populations lead to the full eradication that shrink fast. If it is not fast, the bacterial population shrinking will be reversed and the infection will spread more. This is another signature of the stochastic effects in the bacterial clearance dynamics.

**Figure 3.**
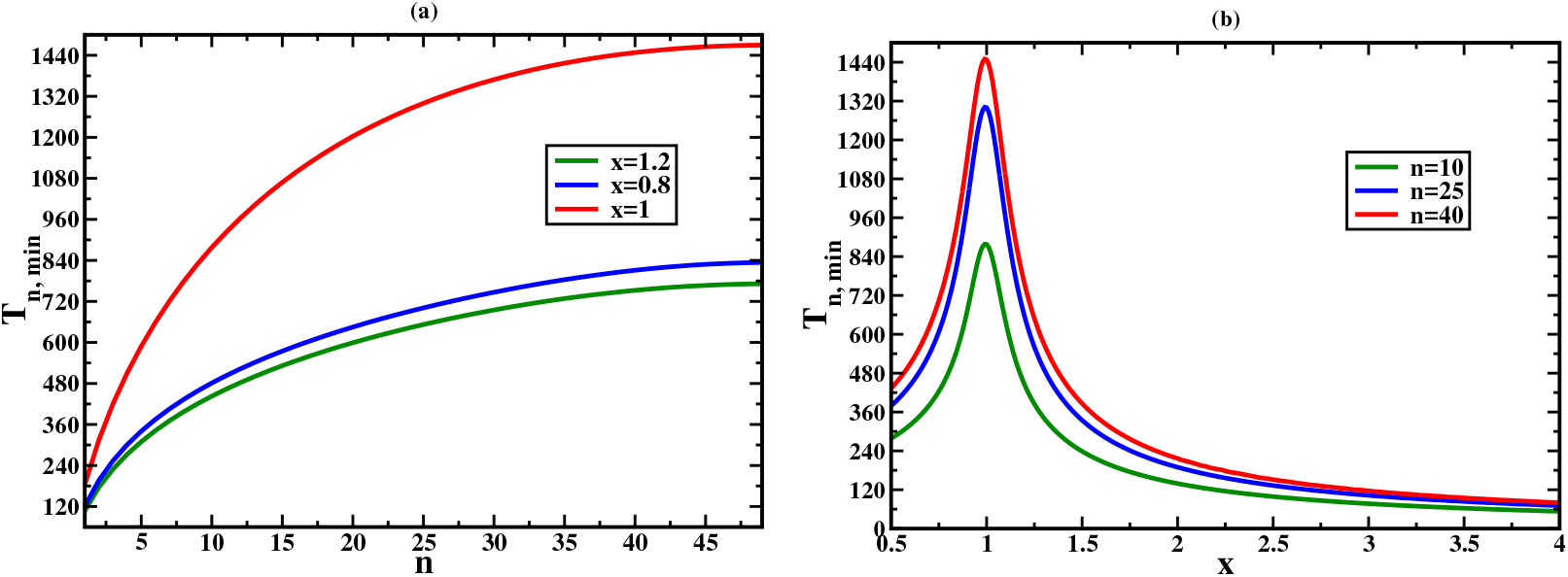
Analytical calculations for the extinction times (in minutes): (a) as a function of the inoculum size for three different values of *x*; and (b) as a function of the parameter *x* for different inoculum sizes (*n* = 10,25, and 40). In all calculations *N* = 50 and λ = 1/60 *min*^−1^ were utilized.

Our analysis of extinction times allows us to reinterpret the meaning of MIC. For *N* → ∞ from (10) we conclude that the extinction time diverges logarithmically for *x* → 1, and it becomes infinite for *x* < 1. This suggests a new more practical definition of MIC (*x* = 1). It is the antibiotic concentration at which the extinction time is maximal (for finite bacterial populations), or it is the antibiotic concentration below which the extinction times diverges (for *N* → ∞). This analysis also suggests that, from the practical point of view, to eliminate the infection it is important to apply the antibiotic concentrations that significantly differ from MIC to avoid the slowdown in the dynamics.

It is interesting to compare our theoretical predictions with experimental measurements of stochastic bacterial clearance [9]. In these experiments, the stochastic population dynamics of bacterial exposed to bactericidal drugs have been monitored starting from single *E.coli* bacteria for sub-MIC conditions (*x* = 0.8) and for concentrations above MIC (*x* = 1.2). It was also estimated that the growth rate is λ ≃ 1/100 *min*^−1^. Then using (10) and (11) we predict that for both cases, *x* = 0.8 and *x* = 1.2, the extinction times are close to 200 minutes, which agrees well with these experimental observations.

### 1.0.2 Stochastic clearance in fluctuating environments

Although the mechanisms of the development of antibiotic resistance remain not fully understood, recent studies suggest that random fluctuations of various parameters can stimulate the bacterial tolerance to antibiotic drugs [2, 15]. In bacterial population dynamics, one main source of stochasticity is due to the environmental variations. For example, single cell experiments have shown that the cell cycle duration is subject to random fluctuations. [30, 31]. We can investigate the effect of growth rate fluctuations on the bacterial clearance dynamics using our theoretical first-passage probabilities method. To do so, we introduce a simplest model as shown in figure 4. It is assumed that the infection can spread with two growth rates, λ_1_ and λ_2_, while the death rate *ϕ* is assumed to be the same in both populations. The system can stochastically transition between two different growth regimes with rates *δ* and *γ*: see figure 4. For the sake of simplicity, in calculations we assume that *δ* = *γ*. Similar deterministic models for population dynamics in fluctuating environments have been already discussed [1, 3, 22].

**Figure 4.**
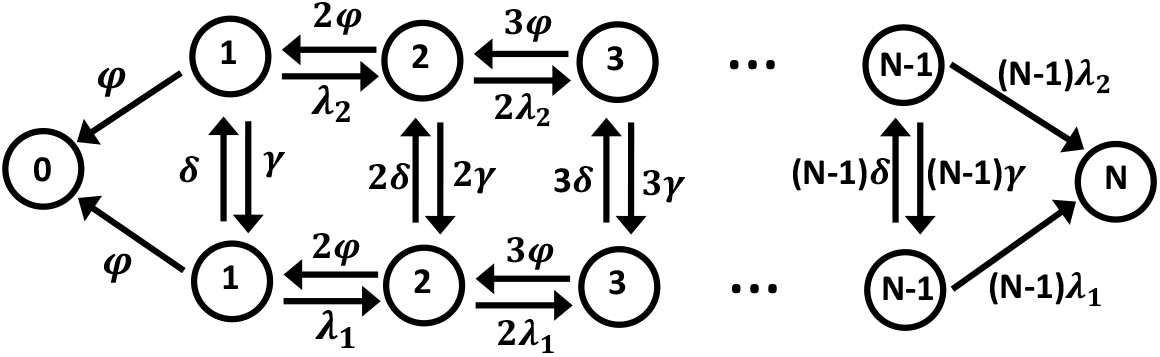
Schematic representation of the model for the clearance of bacteria with fluctuating growth rates. The model comprises two coupled lattices. At each state *n* on lattice 1 (lattice 2), population can jump to state *n* + 1 with growth rate *n*λ_1_ (*n*λ_2_). Death rates are equal along the lattices. Also, *δ* and *γ* are rates to transition between lattices.

In this model, we define 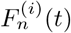 and 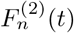 as the probability density functions to clear the system from infection if the bacterial population starts with *n* cells while growing with the rate λ_1_ or λ_2_, respectively. The temporal evolution of these probability functions is governed by the following backward master equations:

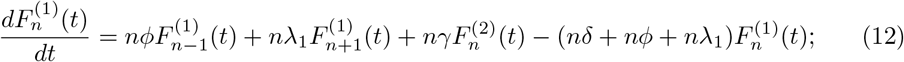

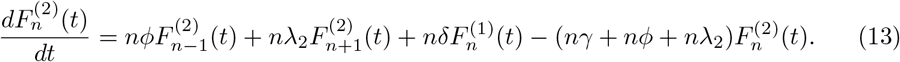

In general, it is difficult to obtain full analytical solution for this problem for arbitrary *N*. However, exact solutions for simple cases with *N* = 2 and *N* = 3 can be derived (see appendix III for details).

To better understand the effects of fluctuation on the dynamics of clearance, it is convenient to compare the fluctuating growth model (rates λ_1_ and λ_2_) presented in figure 4 with a single growth-rate model with 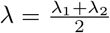 presented in figure 1. Since the average growth rates in both cases are the same, the possible differences in the dynamics properties for bacterial clearance are coming from the fluctuations. To quantify this effect, we define a function 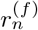 as the ratio of the extinction probabilities predicted by the fluctuating-growth model and by the single growth-rate model:

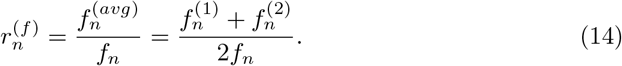

Similarly, one can define a function 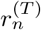 for the ratio of extinction times:

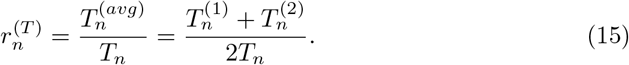

If 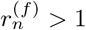 then it means that fluctuations increase the extinction probability, while 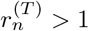 indicates that fluctuations increase the extinction times.

As shown in appendix III, the fluctuating growth-rate model has been solved exactly to evaluate extinction probabilities and extinction times for *N* = 3, and the results are presented in figure 5. It is found that 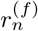 is always larger than one (see figure 5a), which indicates that in the bacterial population with fluctuations in the growth rate the probability of eradication of infection is always larger than in the single-growth population. The effect is stronger for not very large antibiotic concentrations and for slow transitions between two growth regimes. It can be argued that switching transitions open new pathways for eradication of the bacteria, and this should increase the extinction probability. At the same time, increasing the amplitude of the switching transition rates leads to an effective equilibrium single growth rate regime with the growth rate given by the average between two dynamic regimes, and this clearly does not increase the extinction probability.

**Figure 5.**
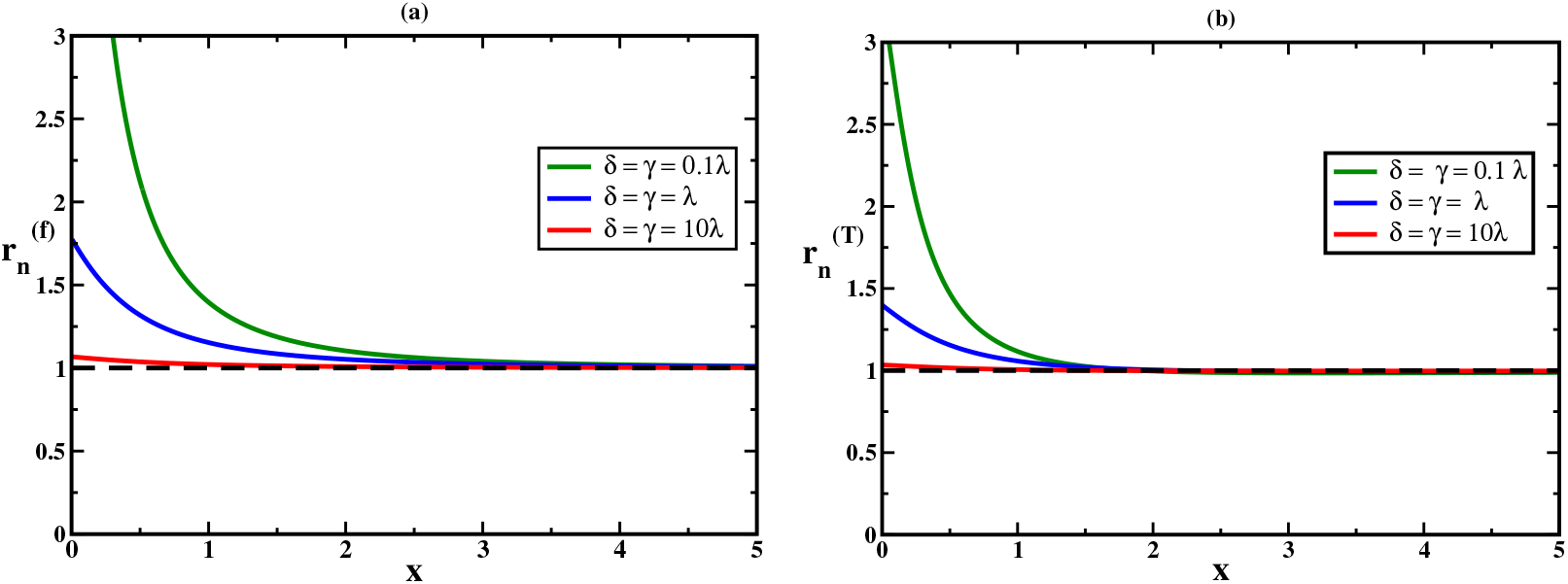
Analytical calculations of dynamic properties for *N* = 3 model. (a) The ratio of the extinction probabilities as a function of *x* for three different values of the transition rates *γ*. (b) The ratio of the extinction times as a function of *x* for three different values of transition rates *γ*. In all calculations, *n* = 2, 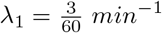, 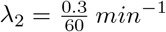 and 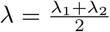 were utilized.

The figure 5b presents the ratio of extinction times, and our theory predicts that 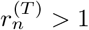, i.e., fluctuations in the growth rates unexpectedly slow down the bacterial clearance dynamics, in contrast to expectations from the extinction probabilities. The effect is stronger for not very large antibiotic concentrations and it disappears for *x* → ∞. It is also strong for weak fluctuation rates between two growth regimes. This surprising result can be explained by noting that due to weak transition rates the system can be effectively trapped in the regime with smaller death rates, and this should slow down the bacterial clearance dynamics.

More realistic situations of bacterial population dynamics require to consider systems with large N. Because analytical calculations cannot be done for these cases, we explored Monte Carlo computer simulations to evaluate the dynamic properties of stochastic bacterial clearance. The results are presented in figure 6. One can see that for relatively small antibiotic concentrations (*x* < 1) the fluctuations in the growth rate increase the extinction probability (figure 6a). In this case, which is generally unfavorable for eradication of infection, opening new pathways should help to clear the infection. This is because the system can spend half of the time in the dynamic regimes with smaller death rates, which helps to fight the infection better. However, the situation changes for large antibiotic concentrations (*x* > 1), when the fluctuations decrease the extinction probability. In this case, due to switching transitions the system spends half of the time in the dynamic regime where it is more difficult to eradicate the infection.

**Figure 6.**
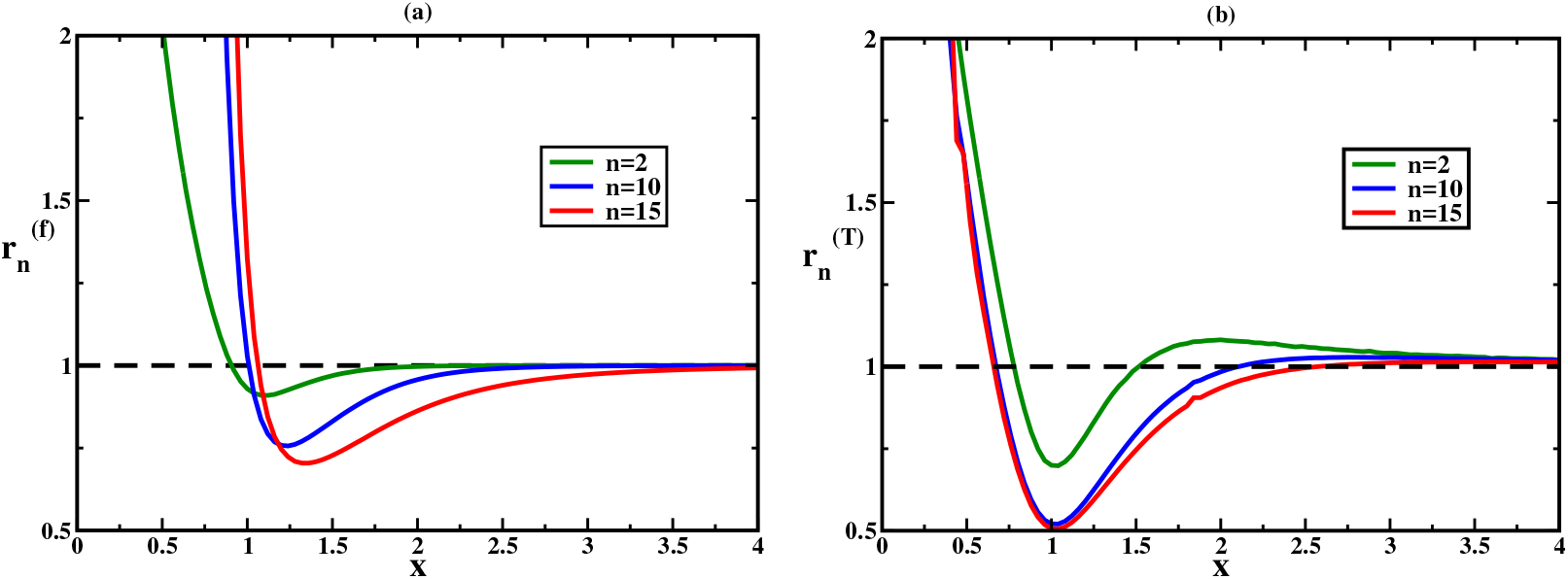
Predictions of Monte-Carlo computer simulations for the fluctuating growth-rate and single growth-rate models. (a) The ratio of extinction probabilities as a function of *x* for three different values of *n*; and (b) the ratio of extinction times as a function of *x* for three different values of *n*. In simulations the following parameters were utilized: *N* = 20, 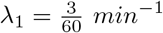, 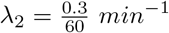, 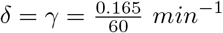.

More complex picture is observed when we analyze the ratio of extinction times: see figure 6b. It is found that for small antibiotic concentrations and for very large antibiotic concentrations the fluctuations in the growth rates lead to slower bacterial clearance dynamics. Only for intermediate antibiotic concentrations around MIC (*x* ∼ 1) fluctuations might accelerate the removal of infection. Apparently, opening new pathways for *x* < 1 and *x* ≫ 1 regions lowers the drive to eradicate the infection because the system spends more time in switchings between different dynamic regimes and not in shrinking of the bacterial populations.

Analyzing the dynamic properties of the fluctuating growth-rate model, two important observations can be made. First, the extinction probability and extinction time generally do not correlate with each other when the system experience fluctuations between different growth regimes. Second, turning on the fluctuations in the growth rates of bacteria can significantly increase the tolerance to antibiotic drugs for large range of parameters. It seems reasonable to speculate that bacteria might explore this option in fighting against antibiotics.

We theoretically investigated the clearance of bacterial population under the effect of antibiotic drugs by concentrating on stochastic aspects of this process. To understand better the mechanisms of eradication of infection, a method of first-passage probabilities is introduced. This allows us to obtain a comprehensive description of bacterial clearance dynamics. Two important dynamic features, extinction probabilities and extinction times, are explicitly calculated. We also clarified the physical meaning of MIC in the systems where the stochasticity is more relevant. Furthermore, using our method we investigated the effect of fluctuations in the growth rates on the bacterial populations dynamics, and we find that these random fluctuations affect differently extinction probabilities and extinction times. For the single growth-rate model, our analysis show that extinction probabilities strongly depend on the antibiotic concentration, the inoculum size and the distance to the death state *N*. But the stochastic effects show up in observations that even for concentrations above MIC the extinction probabilities are not equal to one, while for concentrations below MIC the extinction probabilities are not equal to zero. More complex behavior is observed for extinction times. For finite-size bacterial populations, the extinction times show non-monotonic dependence on the antibiotic concentrations with the maximum at MIC. The unexpected acceleration in the eradication of infection for concentrations below MIC is explained by the fact that the successful events, which are rare at these conditions, must proceed very fast. For infinitely large bacterial populations, our calculations show that the extinction times increase with lowering of antibiotic concentration and diverge for MIC and sub-MIC concentrations. These properties of extinction times provide an additional way of defining the conditions corresponding to MIC.

By introducing a stochastic model in which bacteria can randomly switch between two growth rates we investigated the effect of environment fluctuations in the bacterial clearance dynamics. Our analytical and computer simulations results predict that these switchings increase the extinction probabilities for low antibiotic concentrations, and decrease them for high antibiotic concentrations. However, the effect of fluctuations in the growth rates on extinction times is more complex. With the exception of the intermediate concentrations around MIC, random switchings slow down the bacterial clearance dynamics.

Our calculations lead to several important conclusions. Extinction probabilities and extinction times generally do not correlate with each other, so it is dangerous to make predictions on bacterial population dynamics by considering only the extinction probabilities as typically done in the field. There is a significant range of parameters when the fluctuations in the growth rates lead to the overall slowing down in the eradication of the infection. Bacterial response to antibiotics is a complex process, which depends on genetic and environmental factors [24]. Some bacterial strains are difficult to eradicate because their clearance needs a higher levels of antibiotics that are toxic to hosts. Such bacteria are commonly known as antibiotic-resistant. It is a very challenging task to uncover the mechanisms of the development of bacterial resistance. Our results suggest that one of the first steps in the resistance pathway might be turning on the fluctuations in the growth rates, which would give bacteria an extra time to find another means to avoid the effect of antibiotic drugs. Although at this moment, this is just a pure speculation, it will be interesting to investigate this possibility with experimental methods and more advanced theoretical approaches.

Even at concentrations above MIC, some bacteria survive a short-term exposure to antibiotics before being affected by it. This ability of bacterial population is known as tolerance [6]. In contrast to resistance, which is quantified by the MIC, tolerance is poorly characterized. The most commonly used approach for quantifying tolerance is the measurement of time-kill curves [18]. Recently, a new metric for bacterial tolerance has been introduced [5]. This new metric, known as the minimum duration for killing 99% of the population, *MDK*_99_, can be evaluated by statistical analysis of measurements. Our theoretical method provides the extinction time as a new measure of bacterial tolerance. The advantage of this approach is that it takes into account the stochastic features of the population dynamics and it gives the average dynamic property of the bacterial clearance, which might be much more useful for practical applications.

## Acknowledgments

The work was supported by the Welch Foundation (C-1559), by the NSF (CHE-1664218), and by the Center for Theoretical Biological Physics sponsored by the NSF (PHY-1427654).

## Appendix

### I. Exact solution for the single-growth rate model

In this appendix, we present the details for calculations of the extinction probability and the extinction time. As given in the main text, the temporal evolution of the first-passage probability function is governed by the following backward master equation [27]:

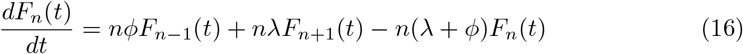

Introducing the Laplace transform of this probability density function, 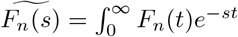, we obtain:

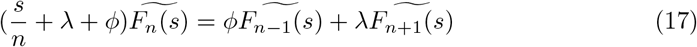

To solve this recurrence relation, it is convenient to write the following expansion:

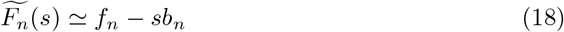

Then 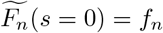 yields the extinction probability. To proceed further we substitute (18) into (17):

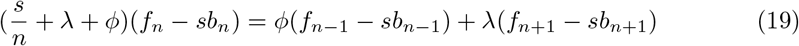

Rearranging terms yields:

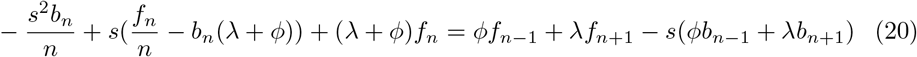

Equating coefficients of *s* on both sides yields two equation recurrence relations:

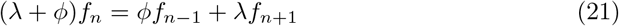

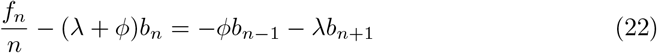

Equation (21) can be simplified as:

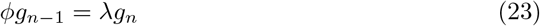

where,

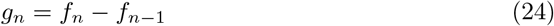

Solution of (24) is given by:

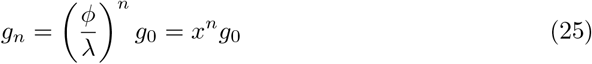

where 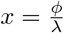. To find constant *g*_0_, we perform summation over equation (24):

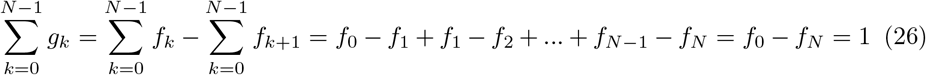

Combining (25) and (26) yields:

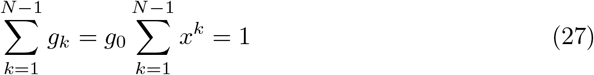

Then, *g*_0_ is given by:

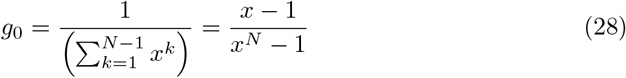

Therefore:

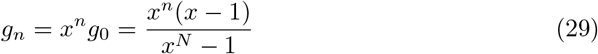

Now using (24), we obtain the extinction probability:

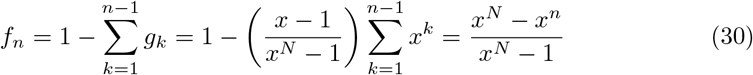

To calculate *b_n_*, we use equation (22)

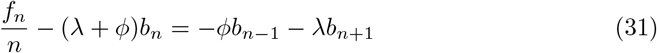

This recurrence relations can be simplified as:

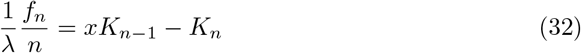

where

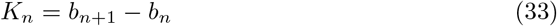

It can be shown that the solution of equation (32) is given by:

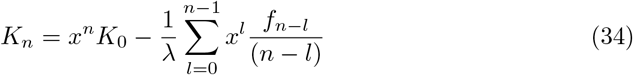

It is convenient to rewrite the summation in the following form:

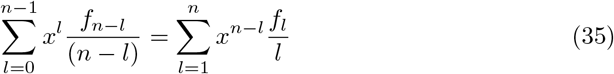

Solution of the recurrence relation *K_n_* = *b_n_*+1 − *b_n_* takes the form:

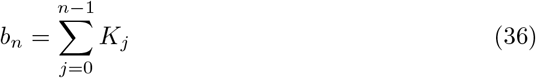

Using boundary condition, we obtain 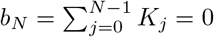. To calculate constant *K*_0_, we perform summation over (34):

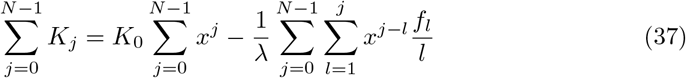

Thus, *K*_0_ is given by:

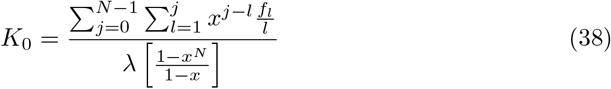

Finally, combining (34) and (36) yields *b_n_*:

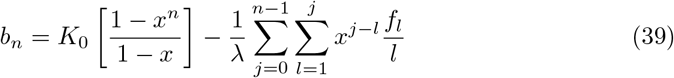

Having determined *f_n_* and *b_n_*, we can now obtain the expression for the extinction time,

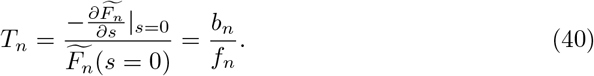

Using (30) and (39), we have

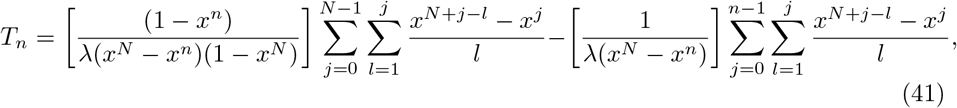

which can be further simplified into

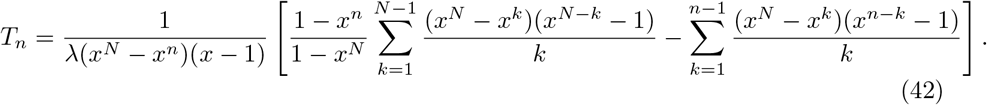

When *x* = 1, this expressions yields

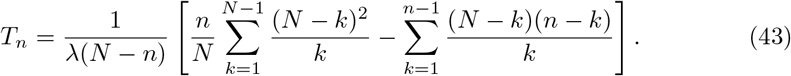

In the case of *x* > 1 and *N* → ∞, it can be shown that

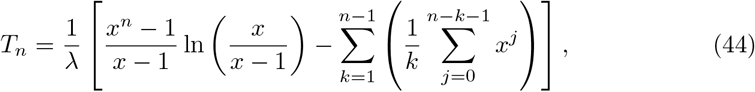

while for *x* → 0 we have

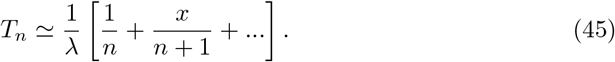

### II. Calculation of saturation point for extinction probability

Since the extinction probability versus *x* follows a logistic sigmoid curve, we can define a saturation value of *x* for which the extinction probability saturates to higher values. There is not a unique way for definition of this saturation point. Here we use a simple definition presented in ([8, 23]). In the simplest approximation, the saturation point is the value of *x* at which the straight line passing through the midpoint (*x* = 1), and with a slope equal to the first derivative of the extinction probability at this point, intersects with *f_n_* = 1. We start by taking derivative of the extinction probability 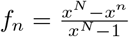 with respect to *x*.

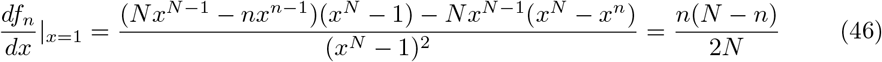

Using this derivate value and coordinate of the midpoint (*x* = 1 and *f_n_* = 1/2), we can obtain the equation of the straight line passing from the midpoint. The equation of line is *y* = *ax* + *b* where 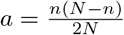. After some algebra we obtain

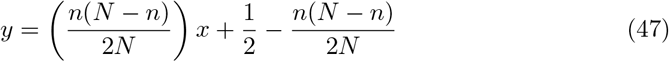

Solution of this equation at *y* = 1 yields the saturation point:

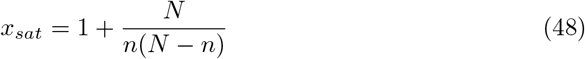

This method only provides a first-order approximation for the saturation point. This approximation can be improved by evaluating higher order (second, third, or fourth) derivatives of *f_n_*. In this case, the straight line passes through the point at which higher derivatives are zero.

### III. Exact solution for the coupled-parallel lattice model

It is difficult to obtain a general analytical solution for equations (12). However, for the small population numbers the exact solution can be derived. In the following we present the details of our calculations for *N* = 3 model.

Schematic of the coupled -parallel mode is shown in (Appendix III - Figure 1). Dynamics of this model is governed by following backward master equations:

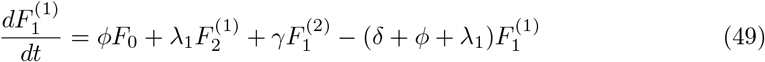

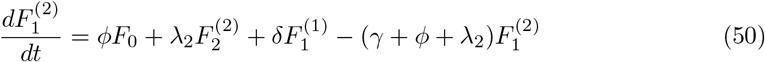

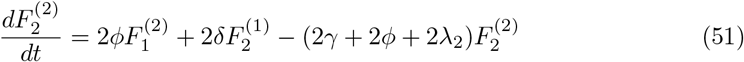

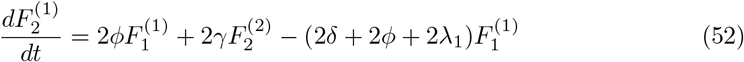

**Appendix III - Figure 1.**
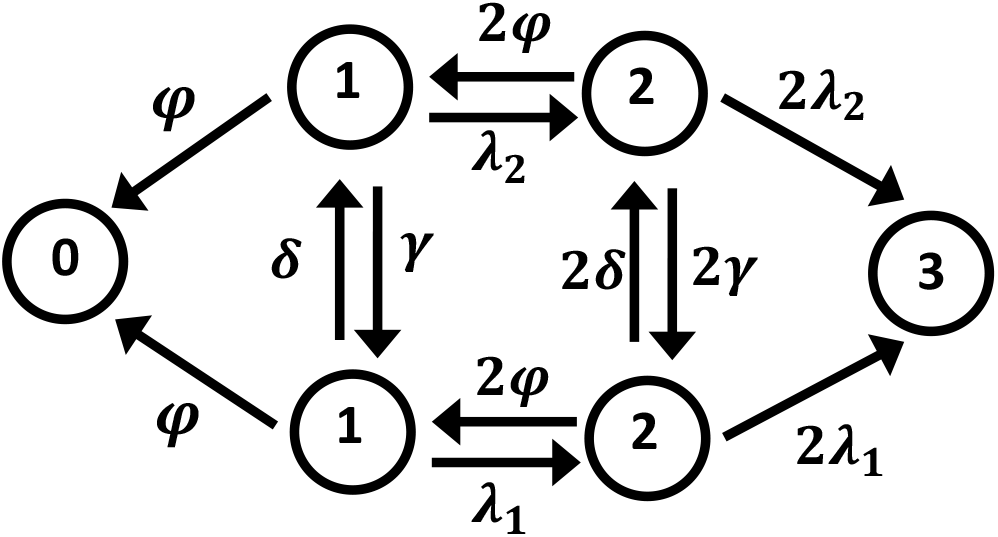
Schematic representation of the fluctuating growth rate model for *N* = 3.

Performing the Laplace transform, we obtain:

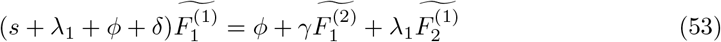

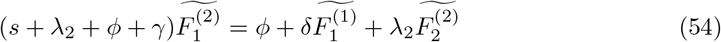

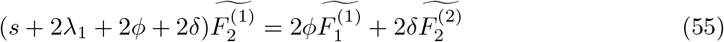

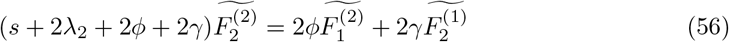

Solving this system of four equations and four unknowns, yields 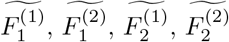. Expanding these functions in terms of *s* yields the extinction probabilities,

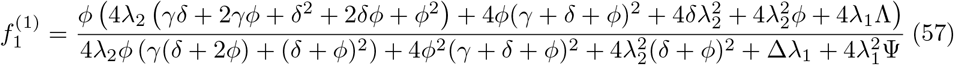

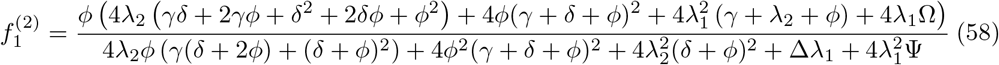

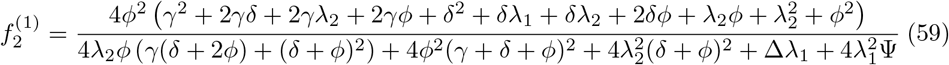

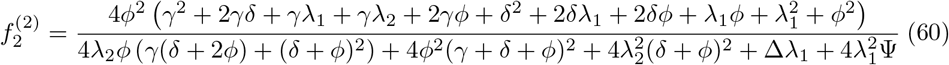

and, the extinction times

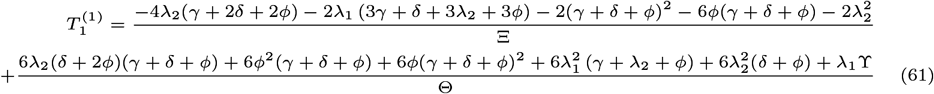

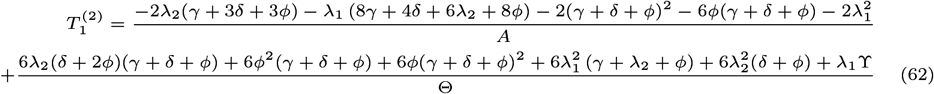

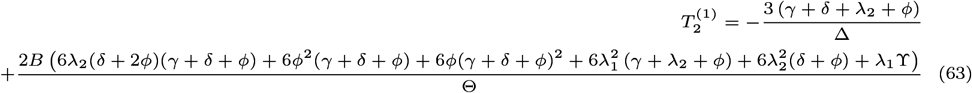

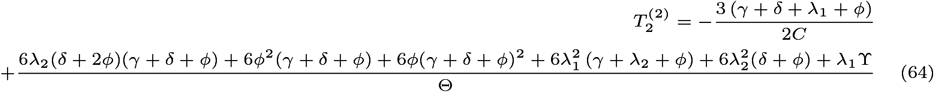

where parameters Ψ, Δ, Λ, Θ, ϒ, Ξ, Ω, *A*, *B*, and *C* are given by:

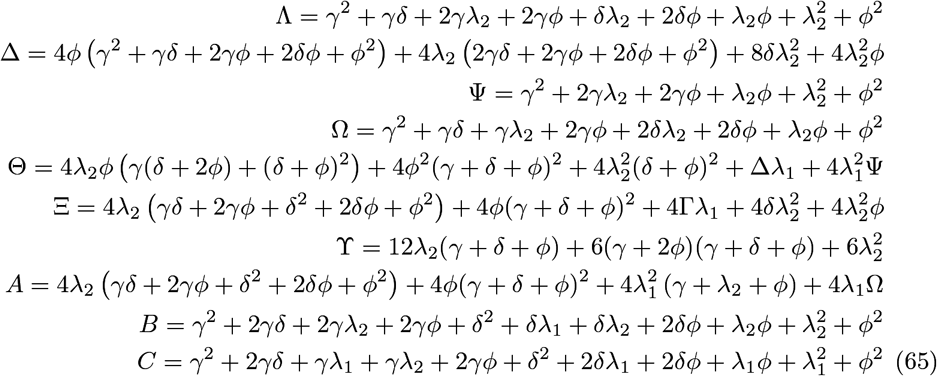

## References

1. M. Acar, J. T. Mettetal, and A. Van Oudenaarden. Stochastic switching as a survival strategy in fluctuating environments. Nature genetics, 40(4):471, 2008.

2. R. Allen and B. Waclaw. Antibiotic resistance: a physicist’s view. Physical biology, 13(4):045001, 2016.

3. N. Q. Balaban, J. Merrin, R. Chait, L. Kowalik, and S. Leibler. Bacterial persistence as a phenotypic switch. Science, 305(5690):1622–1625, 2004.

4. R. Bayston, B. Nuradeen, W. Ashraf, and B. J. Freeman. Antibiotics for the eradication of propionibacterium acnes biofilms in surgical infection. Journal of Antimicrobial Chemotherapy, 60(6):1298–1301, 2007.

5. A. Brauner, O. Fridman, O. Gefen, and N. Q. Balaban. Distinguishing between resistance, tolerance and persistence to antibiotic treatment. Nature Reviews Microbiology, 14(5):320, 2016.

6. A. Brauner, N. Shoresh, O. Fridman, and N. Q. Balaban. An experimental framework for quantifying bacterial tolerance. Biophysical Journal, 112(12):2664–2671, 2017.

7. B. D. Brooks and A. E. Brooks. Therapeutic strategies to combat antibiotic resistance. Advanced drug delivery reviews, 78:14–27, 2014.

8. H. I. Chen and K.-C. Chang. Assessment of threshold and saturation pressure in the baroreflex function curve: a new mathematical analysis. The Japanese journal of physiology, 41(6):861–877, 1991.

9. J. Coates, B. R. Park, D. Le, E. Şimşek, W. Chaudhry, and M. Kim. Antibiotic-induced population fluctuations and stochastic clearance of bacteria. Elife, 7:e32976, 2018.

10. D. Czock, C. Markert, B. Hartman, and F. Keller. Pharmacokinetics and pharmacodynamics of antimicrobial drugs. Expert opinion on drug metabolism & toxicology, 5(5):475–487, 2009.

11. R. Dagan, K. P. Klugman, W. A. Craig, and F. Baquero. Evidence to support the rationale that bacterial eradication in respiratory tract infection is an important aim of antimicrobial therapy. Journal of antimicrobial chemotherapy, 47(2):129–140, 2001.

12. G. V. Doern and S. M. Brecher. The clinical predictive value (or lack thereof) of the results of in vitro antimicrobial susceptibility tests. Journal of Clinical Microbiology, 49(9 Supplement):S11–S14, 2011.

13. M. E. Falagas, G. S. Tansarli, P. I. Rafailidis, A. Kapaskelis, and K. Z. Vardakas. Impact of antibiotic mic on infection outcome in patients with susceptible gram-negative bacteria: a systematic review and meta-analysis. Antimicrobial agents and chemotherapy, 56(8):4214–4222, 2012.

14. B. E. Ferro, J. van Ingen, M. Wattenberg, D. van Soolingen, and J. W. Mouton. Time–kill kinetics of antibiotics active against rapidly growing mycobacteria. Journal of Antimicrobial Chemotherapy, 70(3):811–817, 2014.

15. O. Fridman, A. Goldberg, I. Ronin, N. Shoresh, and N. Q. Balaban. Optimization of lag time underlies antibiotic tolerance in evolved bacterial populations. Nature, 513(7518):418, 2014.

16. P. Greulich, M. Scott, M. R. Evans, and R. J. Allen. Growth-dependent bacterial susceptibility to ribosome-targeting antibiotics. Molecular systems biology, 11(3):796, 2015.

17. E. Gullberg, S. Cao, O. G. Berg, C. Ilbäck, L. Sandegren, D. Hughes, and D. I. Andersson. Selection of resistant bacteria at very low antibiotic concentrations. PLoS pathogens, 7(7):e1002158, 2011.

18. S. Handwerger and A. Tomasz. Antibiotic tolerance among clinical isolates of bacteria. Annual review of pharmacology and toxicology, 25(1):349–380, 1985.

19. R. M. Jones, M. Nicas, A. E. Hubbard, and A. L. Reingold. The infectious dose of coxiella burnetii (q fever). Applied Biosafety, 11(1):32–41, 2006.

20. M. A. Kohanski, M. A. DePristo, and J. J. Collins. Sublethal antibiotic treatment leads to multidrug resistance via radical-induced mutagenesis. Molecular cell, 37(3):311–320, 2010.

21. A. B. Kolomeisky. Motor Proteins and Molecular Motors. CRC Press, 2015.

22. E. L. Kussell, R. Kishony, N. Q. Balaban, and S. Leibler. Bacterial persistence: a model of survival in changing environments. Genetics, 2005.

23. L. M. McDowall and R. A. Dampney. Calculation of threshold and saturation points of sigmoidal baroreflex function curves. American Journal of Physiology-Heart and Circulatory Physiology, 291(4):H2003–H2007, 2006.

24. K. Mitosch and T. Bollenbach. Bacterial responses to antibiotics and their combinations. Environmental microbiology reports, 6(6):545–557, 2014.

25. E. I. Nielsen, O. Cars, and L. E. Friberg. Predicting in vitro antibacterial efficacy across experimental designs with a semi-mechanistic pkpd model. Antimicrobial agents and chemotherapy, 2011.

26. J. O’Neill. Tackling drug-resistant infections globally: Final report and recommendations-the review on antimicrobial resistance. Date of access, 16(09), 2016.

27. S. Redner. A guide to first-passage processes. Cambridge University Press, 2001.

28. R. R. Regoes, C. Wiuff, R. M. Zappala, K. N. Garner, F. Baquero, and B. R. Levin. Pharmacodynamic functions: a multiparameter approach to the design of antibiotic treatment regimens. Antimicrobial agents and chemotherapy, 48(10):3670–3676, 2004.

29. L. B. Reller, M. Weinstein, J. H. Jorgensen, and M. J. Ferraro. Antimicrobial susceptibility testing: a review of general principles and contemporary practices. Clinical infectious diseases, 49(11):1749–1755, 2009.

30. N. Rochman, F. Si, and S. X. Sun. To grow is not enough: impact of noise on cell environmental response and fitness. Integrative Biology, 8(10):1030–1039, 2016.

31. E. B. Stukalin, I. Aifuwa, J. S. Kim, D. Wirtz, and S. X. Sun. Age-dependent stochastic models for understanding population fluctuations in continuously cultured cells. Journal of The Royal Society Interface, 10(85):20130325, 2013.

32. T. Tomita, Y. Fukuda, K. Tamura, J. Tanaka, N. Hida, T. Kosaka, K. Hori, T. Sakagami, M. Satomi, and T. Shimoyama. Successful eradication of helicobacter pylori prevents relapse of peptic ulcer disease. Alimentary pharmacology & therapeutics, 16:204–209, 2002.

33. N. G. Van Kampen. Stochastic processes in physics and chemistry, volume 1. Elsevier, 1992.

34. W. Weidner, M. Ludwig, E. Brähler, and H. G. Schiefer. Outcome of antibiotic therapy with ciprofloxacin in chronic bacterial prostatitis. Drugs, 58(2):103–106, 1999.

35. R. Wilson, S. Sethi, A. Anzueto, and M. Miravitlles. Antibiotics for treatment and prevention of exacerbations of chronic obstructive pulmonary disease. Journal of Infection, 67(6):497–515, 2013.

